# Using digital organisms to investigate the effect of whole genome duplication in (artificial) evolution

**DOI:** 10.1101/521112

**Authors:** Yao Yao, Lorenzo Carretero-Paulet, Yves Van de Peer

## Abstract

The potential role of whole genome duplication (WGD) in evolution is controversial. Whereas some view WGD mainly as detrimental and an evolutionary ‘dead end’, there is growing evidence that the long-term establishment of polyploidy might be linked to environmental change, stressful conditions, or periods of extinction. However, despite much research, the mechanistic underpinnings of why and how polyploids might be able to outcompete non-polyploids at times of environmental upheaval remain indefinable. Here, we improved our recently developed bio-inspired framework, combining an artificial genome with an agent-based system, to form a population of so-called Digital Organisms (DOs), to examine the impact of WGD on evolution under different environmental scenarios mimicking extinction events of varying strength and frequency. We found that, under stable environments, DOs with non-duplicated genomes formed the majority, if not all, of the population, whereas the numbers of DOs with duplicated genomes increased under dramatically challenging environments. After tracking the evolutionary trajectories of individual artificial genomes in terms of sequence and encoded gene regulatory networks (GRNs), we propose that increased complexity, modularity, and redundancy of duplicated GRNs might provide DOs with increased adaptive potential under extinction events, while ensuring mutational robustness of the whole GRN. Our results confirm the usefulness of our computational simulation in studying the role of WGD in evolution and adaptation, helping to overcome the traditional limitations of evolution experiments with model organisms, and provide some additional insights into how genome duplication might help organisms to compete for novel niches and survive ecological turmoil.

## Introduction

Polyploidy, or the duplication of entire genomes, is a common phenomenon in the evolutionary history of many eukaryotic organisms, especially plants. Whole genome duplications (WGDs) have often been linked to speciation and species diversification, increase in biological complexity, alleviating the effects caused by extinction events, and an overall increased environmental and mutational robustness (1–3). Nevertheless, although the prevalence of WGDs has now been firmly established, the effects and significance of WGDs for evolution remain vividly discussed. Whereas some regard WGD mainly as an evolutionary dead end, others see WGD primarily as an opportunity and source of evolutionary innovation. In support of the former, there is the observation that recent WGDs seem to outnumber established ancient polyploidy events by several orders of magnitude (4–6). This relative paucity of anciently established WGDs seems to suggest that many WGDs have not survived on the long run, which may be due to the well-known detrimental effects of polyploidy, caused by minority cytotype exclusion, genomic instability, mitotic and meiotic abnormalities, or epigenetics changes (7).

On the other hand, we do observe ‘ancient’ organisms that underwent and survived WGDs, so that their descendants have outcompeted their diploid progenitors and bear traces of the duplication in their genomes. This is the case for several major eukaryote lineages such as vertebrates (8, 9), fishes (10), ciliate protozoans (11), hemiascomycetous yeasts (12, 13), and particularly plants (14–16). This observation, together with earlier work on specific adaptations of polyploids, led some to conclude that WGDs might provide an adaptive advantage particularly under unstable, challenging and stressful environments or during periods of environmental upheaval (6, 17). This hypothesis is supported, among others, by the wave of successful WGDs that seems to have occurred around the Cretaceous-Paleogene extinction event or K-Pg boundary (18–21).

Although there is increasing support for a correlation between polyploid establishment and environmental challenge (6, 17), the molecular evolutionary mechanisms behind this remain vague. Ideally, to comprehensively investigate the effects of WGDs for evolution and study the link between adaptation and polyploidy, we need to collect detailed evolutionary trajectory data of organisms with and without WGDs (e.g., the complete mutational landscape, dynamic gene expression profiling data, real-time adaptation data, etc.) under different environmental contexts. However, although there have been some very interesting and successful attempts, collecting these data from real biological evolution experiments is difficult and bound by experimental limitations (22). Computational approaches, notwithstanding many other limitations, have the advantage that they can collect at least some of such evolutionary trajectory data and have already shown great potential in investigating the potential adaptive role of WGD. For example, computational evolutionary modeling of populations of so-called ‘virtual cells’ has been used to study the adaptive potential of polyploidy in times of drastic and enduring environmental change (23). Although WGD was established only in a minority of lineages, polyploids were significantly more successful at adaptation (and readaptation), and therefore WGD seemed, at least in some cases, a powerful mechanism to cope with environmental challenges. However, as far as we understand, the model of Cuypers et al. (23) does not allow modeling the connection of the gene regulatory network (GRN) of these virtual cells with a particular environment.

We have recently published a computational framework aimed at mimicking biological evolution (24–26). Our framework is based on so-called ‘digital organisms’ (DOs), which have their own genomes replicating under specific mutation and WGD rates. Furthermore, the expression of every gene in the genome is regulated by a set of environmental signals, or sensors, and the orchestrated activity of regulatory genes defined in the genome. The overall gene expression pattern of the entire gene sets in the genome defines the GRN of the DO, which in turn determines its behavior, defined by a set of actuators. The strengths and advantages of our bio-inspired agent-based modeling approach have already been tested under a relatively simple setup, such as emergent swarm behavior and overall enhanced adaptation in a changing environment (24, 25).

Here, we expand our previously developed framework to study the effects of polyploidy in a challenging environment. By considering the adaptation of each individual DO as an interactive and developmental process in evolution, we have a better chance to observe when and under which circumstances polyploidy can pose a potential selective advantage to the organism or population as a whole. Because the effects and consequences of polyploidy are complex and probably also often lineage-specific, we would like to stress that in the present study, we only focus on how polyploidy or WGDs might affect the evolution of GRNs and what might be some of the major constraints on the evolution of GRNs, and hence adaptation of their hosts, subsequent to WGDs. We do not study (the probability of) short-term establishment of polyploidy, nor longer-term phenomena such as gene loss or the functional divergence of genes (sub- or neofunctionalization). Also, reproductive effects (e.g. sexual reproduction and recombination) and population-specific features, such as allele frequency changes following WGDs, are not considered here. Although we are of course aware that these are all very important to understand the significance of polyploidy and WGD, we nevertheless hope that, by focusing on certain molecular properties, some aspects of WGDs can be better understood.

By analyzing multiple simulations of populations of clonal DOs under various environmental contexts, we found that DOs with non-duplicated genomes conformed most, if not all, of the population, under stable environments. In contrast, environmental stress or change usually resulted in higher rates of extinction in the population, whereas WGDs seemed to increase the possibility of survival under dramatically challenging environments. Furthermore, tracking mutational changes in populations led us to observe that the GRNs in duplicated genomes tend to show greater flexibility with fewer mutations compared to those in non-duplicated genomes. Our results provide some initial insights into how genome duplication might help organisms to diversify, compete for novel niches, and survive ecological turmoil.

## Results

### Studying the Impact of WGD on the Evolution of DOs Under Different Environmental Conditions

To study the putative effect of WGDs on the evolution of DOs under challenging environmental conditions, we have simulated the evolution of clonal populations of DOs under two different scenarios of food availability and specific rates of WGD and nucleotide substitution. It should be noted that, at every time step and for every simulation, the population size of polyploids and non-polyploids was computed. Although, after food removal, a drop in population size was generally observed, when and how and to what extent the population size drops can greatly vary from simulation to simulation (25). It is inherent to the system that DOs with different genomic makeups and encoded GRNs can have very different responses to the same amount of food removal, which makes it hard to summarize the results, except for evaluating overall success when putting polyploids and non-polyploids in competition. In the first scenario, food was dynamically removed every 60 time steps after the total population (polyploids plus non-polyploids) reached 1000 individuals. Starting from a population of 200 individuals at t_0_, having reached a population size of 1000 indicates that at least part of the population has adapted to the current environment and started to replicate rapidly (24, 25). Under this first scenario, we performed 30 simulations under each of the following four food reduction levels: no food reduction, except for what is found and consumed by the DOs, 50% food reduction, 70% food reduction, and 85% food reduction. After 2000 time steps, we compared the population sizes of polyploids and non-polyploids (Table 1). As can be observed, under no or low food reduction scenarios, at the end of the experiment, most of the population consisted of non-polyploid DOs (genomes). The population size of polyploids increased with the percentage of food removed, becoming bigger than that of non-polyploids only when 70% of the food was removed repeatedly, i.e. every time the total population reached 1000. Taking away 85% of the food resulted in the extinction of the whole population of both polyploids and non-polyploids in all simulations.

**Table 1.** Summary of results from evolution simulation experiments under dynamic and continuous food reduction every time the population has reached 1000 individuals. For every food reduction group, 30 simulations were run 3 times during 2000 time steps.

|  | No food reduction | 50% food reduction | 70% food reduction | 85% food reduction |
| --- | --- | --- | --- | --- |
| Extinction (for all DOs) | 0% | 10% | 16.7% | 100% |
| Polyploid population bigger than non-polyploid population | 10% | 30% | 63.3% | 0% |
| Non-polyploid population bigger than polyploid population | 90% | 60% | 20% | 0% |

In the second scenario, the simulation ran for 1000 time steps but ‘food removal’ was introduced at a fixed time step, namely at time step 300. Because the growth of food throughout the experiment is exponential, introducing food reduction at time step 300 represented an appropriate balance between earlier reduction before polyploid and non-polyploid populations have reached an equilibrium in the total population, and later reduction, which may not allow to remove enough food for re-adaptation or efficient selection to occur. Four different levels of food reduction (0%, 30%, 60% and 90%) were implemented, each with 10 different populations starting with 200 clonal DOs each. The population size of polyploids and non-polyploids was compared at time step 1000 (Table 2). Similarly to what we observed under the previous scenario, at 0 and 30% of food reduction, no polyploid genome survived at the end of the experiment, with some polyploid genomes surviving only under a 60% or more of food reduction. Polyploid populations were able to overtake non-polyploid ones only when significant levels of food (90%) were removed (Table 2).

**Table 2.** Summary of results from evolution simulation experiments under fixed food reduction at time step 300. For every food reduction group, 10 simulations were run during 1000 time steps.

|  | No food reduction | 30% food reduction | 60% food reduction | 90% food reduction |
| --- | --- | --- | --- | --- |
| Extinction (for all DOs) | 0% | 0% | 0% | 20% |
| Polyploid population bigger than non-polyploid population | 0% | 0% | 20% | 60% |
| Non-polyploid population bigger than polyploid population | 100% | 100% | 80% | 20% |

In summary, both simulation scenarios show that, while non-polyploids clearly outcompete polyploids when environmental conditions are stable, when major environmental changes are introduced in the form of drastic reductions of food available, polyploids seem to confer an advantage over non-polyploids. This effect is more noticeable under the first scenario, where the environmental changes are repeated multiple times through recurrent food removal, creating stronger environmental constraints and hence greater needs for multiple re-adaptations (see further).

### Evolution of the mutational landscape of DO genomes following WGD

In order to gain further insights on why and how polyploid DO populations have, in most simulations, a greater chance of surviving extreme environmental changes (measured by the number of surviving DOs at the end of the simulation), we examined the evolution of the average long-term genetic distance (LTGD) and short-term genetic distance (STGD) of ‘adapted’ populations. When referring to an ‘adapted population’, we mean that each DO in the population has an energy level of at least 20,000 units at that particular time step. This ensures that we only reconstruct the evolutionary trajectories from organisms that have had a chance for replication in the population. Fig. 4 summarizes the result for 30 simulations under a 70% food reduction scenario. As can be observed, the average STGD of non-polyploids is significantly higher than the average STGD for polyploids during most time steps (0.1876360 < 0.1325995, p-value < 0.02, two sample *t*-test, Fig. 4*B*), whereas the opposite is true for average LTGD values (2.646769 > 2.215343, p-value < 0.02) (Fig. 4*A*).

**Fig. 1.**
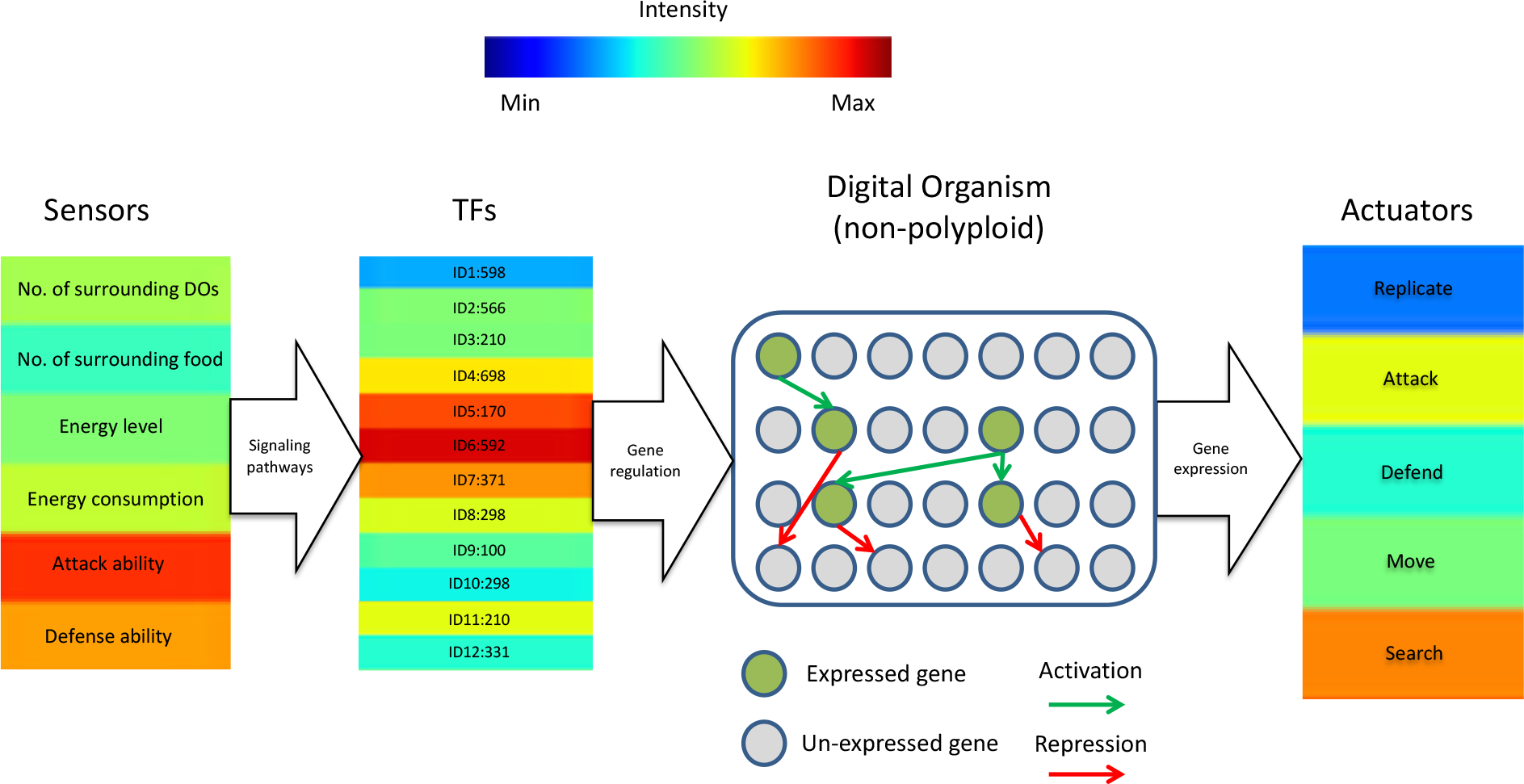
Sensors, actuators and the genome of a digital organism in our evolution simulation computational framework. At every time step in the simulation, the values of the sensors are converted by means of linear equations into a number. This number represents the specific identity number of a particular gene product, resulting in an increase of its corresponding concentration level. The concentration level of all gene products determines the overall behavior of a DO, which is in turn defined by the set of possible actuators, each one encoded by its corresponding structural genes

**Fig. 2.**
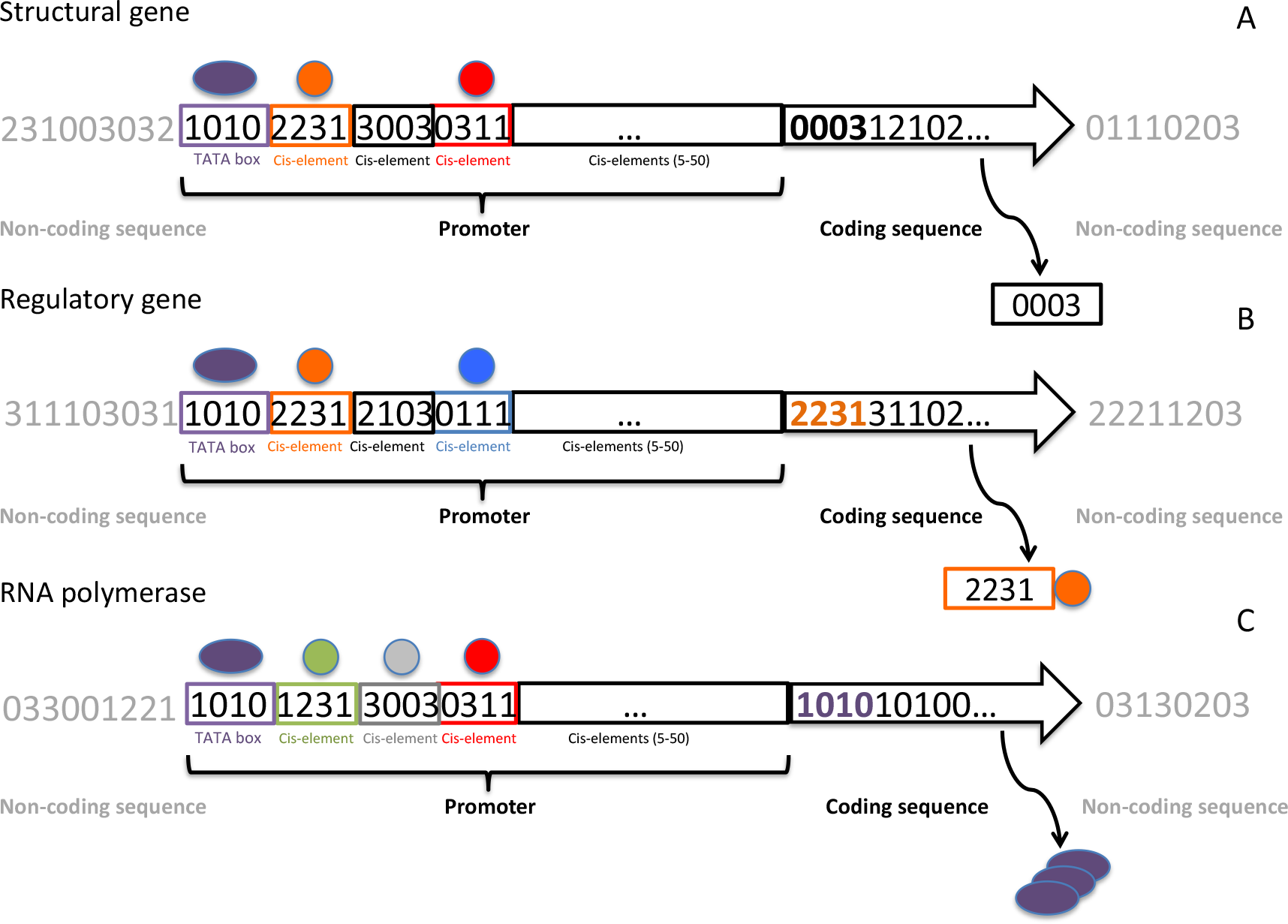
Schematic representation of the three kinds of genes encoded in the genome of a digital organism. Genes in the genome are randomly initialized in every newly built genome. Every gene is defined by a promoter and a coding region. The promoter must start with a TATA box and be followed by a set of four-digit cis-elements numbering 5-50, each of which is bound by a specific kind of TF. The promoter region is followed by the coding region, whose first four digits define the specific kind of gene to be encoded. No interspersed sequence is allowed between any of the components of the gene. (*A*) Structural gene, which defines a specific kind of actuator. (*B*) Regulatory gene, which encodes a specific kind of TF defined by its cis-binding element (i.e., 2231). (C) RNA polymerase gene, which specifically binds to a TATA box (i.e., 1010)

**Fig. 3.**
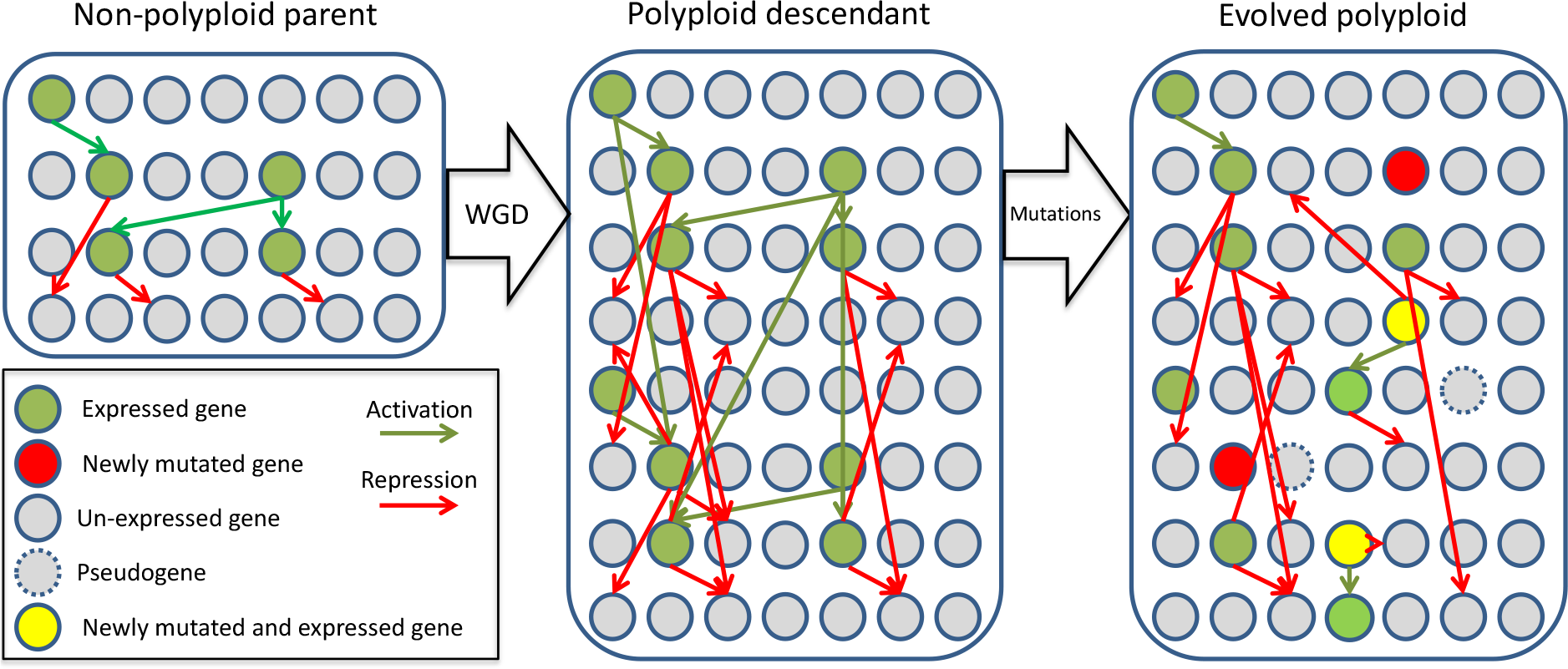
Evolution of a DO under specific WGD and substitution mutation rates. At the beginning of the simulation, the WGD rate is fixed at 40%. Once non-polyploid and polyploid populations sizes have reached an equilibrium, the WGD operator is removed from the simulation. Similarly, at every replication step, a fixed substitution rate (10^−4^ per base) is set

**Fig. 4.**
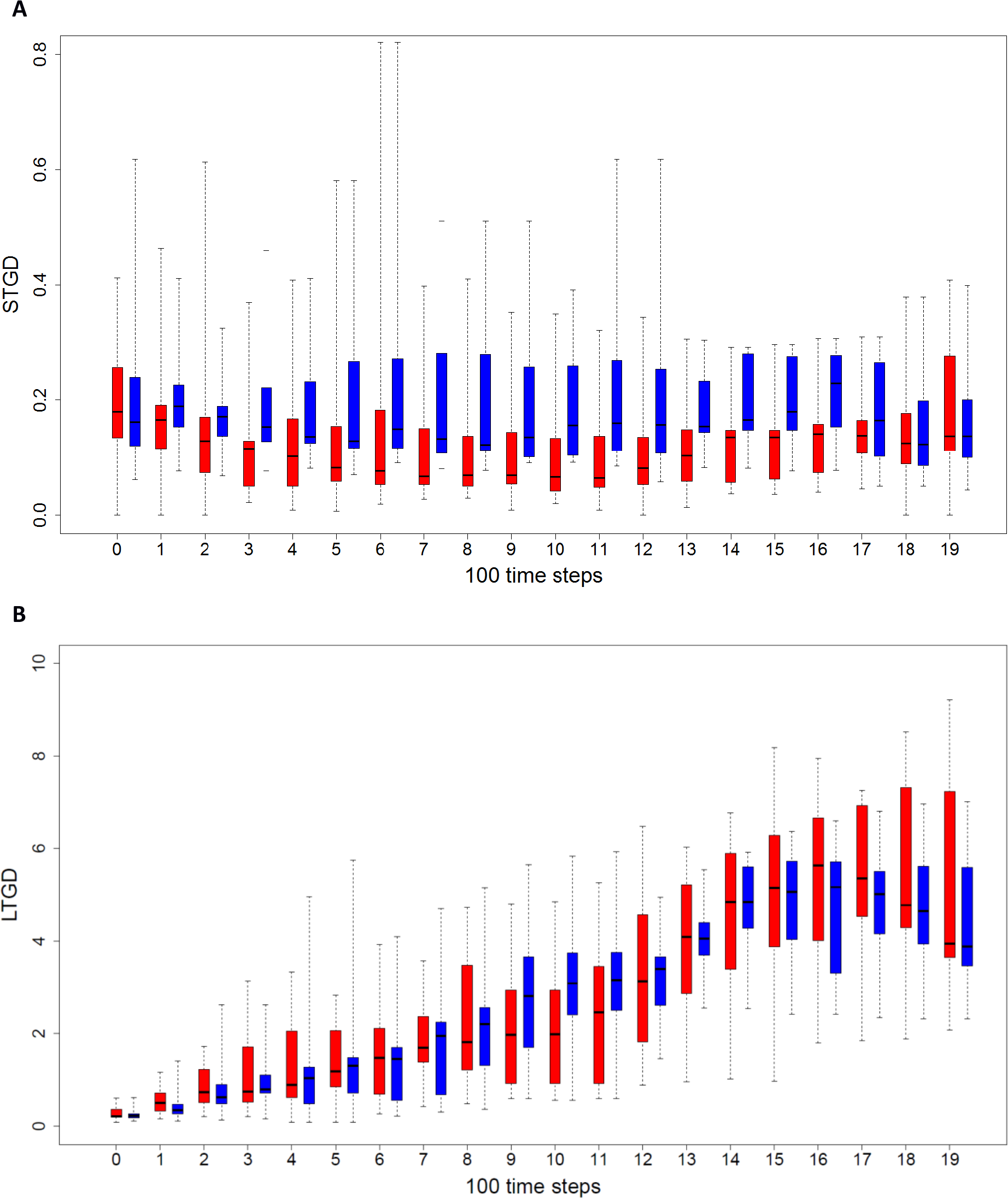
Evolution of short-term genetic distance (*A*, STGD) and long-term genetic distance (*B*, LTGD) across 30 experiments (simulations) under a dynamically changing environment (70% food reduction) Red and blue boxes represent the ‘adapted’ polyploid and non-polyploid populations, respectively. Solid lines in box blots represent the median values of the STGD and LTGD of all simulations, while the box borders correspond to the first and third quartile and the extreme values correspond to the lowest and highest values observed in any of the simulations. See text for details

It should be noted that the substitution rate for all genomes is the same and hence, under the assumption that WGD has no effect on mutational patterns, the average LTGD and STGD should also be the same for both populations. Consequently, the lower values for the STGD in polyploids have to be the result of selection and thus selection on the short-term seems more stringent for polyploids than for non-polyploids, although this does not seem to be the case for long-term evolutionary processes. This would suggest that, with a duplicated genome structure, random substitutions on genes are under more stringent selection, but once selected and inherited, mutations also have more chance to be retained for longer times (see further).

### Evolution of GRNs following a WGD

We also investigated the effect of WGDs on the encoded GRNs by measuring the extent of changes in the concentration levels of its different components (gene products). Because of the limited mobility of DOs, most of the time during our simulated evolution experiments, the surrounding environments could be considered relatively stable, no matter how and when abrupt environmental changes were introduced. Though abrupt food removal may cause a rapid change in the underlying GRN of the DO’s genome (a change in food availability can change the environmental signals resulting in domino effects on the concentration levels of particular TFs), such changes were usually limited in time. From this, we inferred that adapted DOs would maintain a relatively stable GRN most of the time. This assumption provided us with a way to roughly identify the DOs that have the potential for long-term adaptation. DOs with more stable GRNs will be more capable to maintain their adaptive features, in contrast to DOs showing more divergent expression patterns and GRNs. Consequently, we considered the stability of GRN expression patterns (represented by the expression distance (ED), see Methods) as an indicator for adaptive potential, and used this indicator to investigate the effect of WGDs in the long term. For this purpose, we extracted a subpopulation of DOs that had likely established a comparatively stable GRN (ED values lower than 30%, reflecting homeostasis) and reached an at least temporary adaptive state (energy levels above 20,000). The evolutionary trajectories in terms of LTGD between polyploids and non-polyploids in this subpopulation were then compared to that of the total population of DOs in each simulation. For this purpose, we ran 40 additional simulations under a scenario of 70% of dynamic food reductions (Fig. 5). We observed that non-polyploids usually need to accumulate considerably more mutations (hence resulting in a higher LTGD, at least for the adapted subpopulation) to establish novel GRNs that have adapted to the new environment (Fig. *5B*), while for the polyploid subpopulation, the difference was less noticeable. In the current implementation of our framework, we do not know whether this is due to positive selection or selective sweeps in one group of DOs or due to purifying selection in the other, but it is a fact that different regimes of selection are ongoing in polyploids and non-polyploids. This contrasting pattern between polyploids and non-polyploids is more obvious at later steps of the simulation, suggesting that polyploids and non-polyploids have indeed different ways to respond to environmental changes. In other words, although there is still a considerable fraction of non-polyploid DOs in the population, based on great fluctuations of their gene expression (large ED values), we can infer that their encoded GRNs have not (yet) adapted to their changing environment. Therefore, these DOs may survive in the short term, but will have reduced chances of surviving in the long term, probably being eliminated by environmental fluctuations. In any case, based on these observations, we can infer that, for re-adapting under changing environments, non-polyploid genomes need to accumulate more mutations, i.e. need to make bigger jumps in phenotype space (see Discussion) than polyploid ones (Fig. 5).

**Fig. 5.**
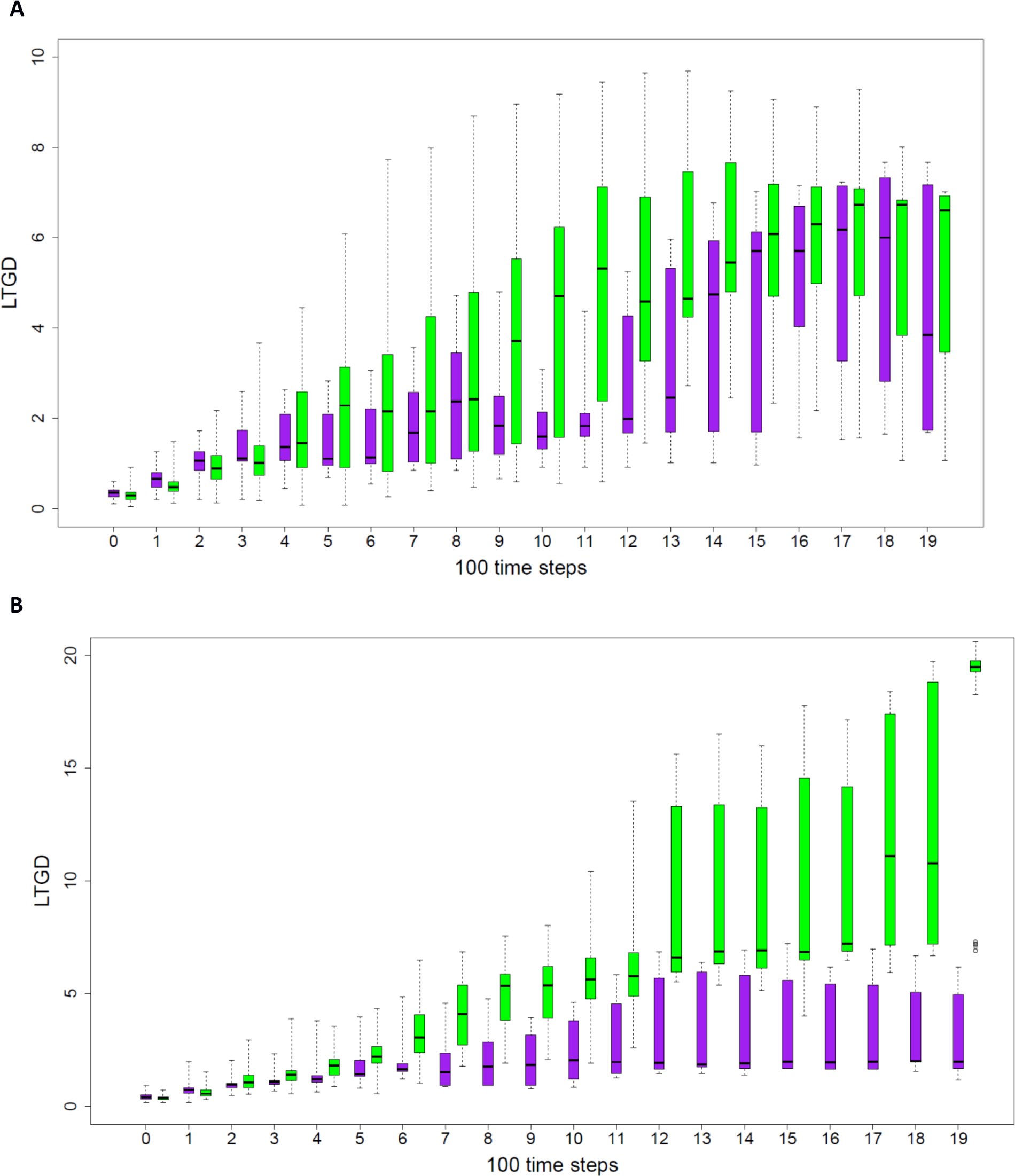
Evolution of the average LTGD of a subpopulation of DOs with stable and adapted GRNs versus all DOs from 40 experiments (simulations) under a dynamically changing environment (70% food reduction) Purple boxes represent the overall population and green boxes represent the population with stable GRNs (i.e. considered adapted). (*A*) polyploid populations; (*B*) non-polyploid populations. Interpretation is as in Fig. 4

## Discussion

With the purpose of investigating whether WGDs may confer an adaptive advantage in the face of drastic environmental changes or extinction events, such as abrupt reductions in available food resources, we have implemented here an improved version of our previously designed bio-inspired and agent-based computational framework (24, 25). Two different environmental scenarios were set: food reduction was either introduced at a fixed time step during the simulation, or dynamically every time the total population reached 1000 individuals. From our simulations, we learned that under stable environments, non-polyploids in general do better than polyploids, as reflected by the low or null fraction of polyploids surviving at the end of the different simulations performed under low or null food reduction. Under both scenarios, the relative fraction of polyploid DOs surviving at the end of the experiment increased with increasing levels of food removal, overcoming the population of non-polyploid DOs when the environmental challenge introduced in terms of food reduction was strong enough. Taken as whole, these results suggest that WGDs are usually maladaptive under stable environments, but may actually confer an adaptive advantage under certain environmental constraints, which seems in agreement with recent ‘real-life’ observations (6, 27–29).

Furthermore, we observed that in DOs with duplicated genomes, fewer mutational changes are allowed to accumulate (as measured by STGD) than in DOs with non-duplicated genomes, suggesting the former are under stronger purifying selection. We have also shown that DOs with non-duplicated genomes need to accumulate many more mutations than polyploid genomes - hence their larger LGTD values - to reach adaptation (using homeostasis as measured by ED and the energy level as proxies). We interpret this as follows: under stable environments, and in ‘well-adapted’ organisms with (relatively) stable GRNs, there is always a certain chance that a random mutation renders the GRN (and consequently its host) less well adapted to that environment. This phenomenon is exacerbated in more complex GRNs. Indeed, the more complex the GRN, the more likely it is that a mutation has a wider impact. With an increasing number of edges and connections in the network, the probability increases that a random mutation in a regulator affects a larger number of genes, in turn increasing the chance of causing major disturbances to the underlying GRN (30–32). In this respect, WGDs increase the number of edges in the network exponentially (Fig. 2). Concretely, while the average GRN for the total population consisted of 272 instances or agents at the start of our simulations, this increased to about 1,700 for DOs with a duplicated genome at the end of the simulations.

A duplicated genome also results in an increase in the modularity of its encoded GRNs, i.e., the occurrence of multiple functional subnetworks or modules formed by highly connected genes that are co-expressed and/or co-regulated by the same set of key regulators, while establishing sparser connections with nodes from different modules (1, 33–38). As a consequence, single substitutions in duplicated genomes have the potential to affect many more key regulators, in turn having a greater impact on the underlying GRN and the resulting phenotypes. Whereas this may be disadvantageous in stable environments, under drastic environmental changes, DOs with duplicated genomes may actually have a better chance to acquire the drastic genomic changes necessary to survive. Furthermore, the modular arrangement of duplicated GRNs may also result in functionally redundant modules, contributing to the genetic or mutational robustness at the GRN level. This facilitates the rewiring of novel functional modules without disturbing the ancestral one, a feature that is expected to be especially advantageous under unstable challenging environments (23, 36). In this respect, our observations are compatible with previous claims that increased modularity results in increased evolvability (32, 39-44). The origin and evolution of modularity seems, at least in some cases, to be linked with repeating elements such as duplicated genes (38, 45, 46). For a population that is already adapted to a certain environment, the increased modularity of the duplicated GRNs could be maladaptive or detrimental because the GRNs are more vulnerable to random mutations and the potential for increased evolvability may have little value or direct benefits. On the other hand, the modular structure of duplicated GRNs might allow polyploids to explore a wider evolutionary landscape, providing short-term increased opportunity to adapt to novel, different, or rapidly changing environments, while redundancy allows the gradual evolution of different functional subnetworks for adapting in the longer term. In partial summary, increased complexity, modularity and functional redundancy following WGD would help to explain the different behavior of polyploids and non-polyploids under stable or challenging environments (6).

Although many other factors may be at play as well, which, for simplicity, are not considered here (but will be addressed elsewhere), the effect of WGD on network complexity, modularity and redundancy could be part of the explanation why also in natural environments, floras in stable environments have lower proportions of polyploids than highly, recently disturbed floras. In a recent study, Oberlander et al. (47) suggested that the hyper diverse South African Cape flora, which has anomalously low proportions of polyploids compared to global levels, might be due to climatic and geological stability, supporting the hypothesis that WGD may be rare in stable environments. On the contrary, there is considerable evidence that polyploidy is much more common in disturbed habitats, or habitats that (have) show(n) environmental turmoil (6, 27, 48). Previously, we have also wondered about the discrepancy between the existence of many polyploids of fairly recent origin and the scant evidence of ancient polyploidy events, certainly within the same evolutionary lineage (6). The paucity of polyploidy events that ‘survive’ and are established in the long term would suggest that polyploidy is usually an evolutionary dead end. Again, this fits with our observation that the increased complexity of the underlying networks may be detrimental (or at least sub-optimal) for species that are well-adapted to their environment, when accumulating (too) many changes. In this respect, it is also noteworthy to mention that during our simulations, rarely two or three WGDs were observed, but never more. Also in real-life organisms, very few examples are known of consecutively established WGDs that have occurred in a short period of time. Some of these presumed exceptions are *Musa acuminata*, *Spirodela polyrhiza*, and *Arabidopsis thaliana* (6). However, at least for *Arabidopsis* (49) and *Spirodela* (50), the number of genes is very low compared to what would be expected following several rounds of WGD, suggesting huge gene loss and fractionation following WGD. In line with our observations made in the current study, quickly losing a large amount of the extra genetic material created through a WGD might have facilitated surviving these major events.

Our observations might also have some broader implications and might help to understand the origin of complex phenotypes and discontinuities in biological evolution. Traditionally, evolution is regarded as progressing at a more or less constant rate, which is linked to the gradual accumulation of small genetic changes over time. This model successfully explains, for instance, the variation between (closely) related species but often fails to explain bigger ‘leaps’ in evolution. However, there is ample recording of ‘novel’ or highly divergent species that seem to have emerged in a relatively short evolutionary time frame and sometimes even representing important evolutionary transitions. Such observations often seem difficult to reconcile with an evolutionary model that is based on the gradual accumulation of small changes, and a huge body of literature has been devoted to this seemingly contradictory phenomenon (51, 52).

As supported by our simulations, and in previous research (24, 25), in a stable environment, gradual evolution can successfully explain the divergence and optimization of evolutionary processes. When, on the other hand, the environment drastically changes, in a short geological time frame, gradual evolution has difficulty to keep up, while new species or life forms, if different enough from the original ones, can successfully occupy the ‘new’ niches that have become available (3). When the environment is rapidly changing, existing species may not have enough time to adapt and will disappear. WGD provides a way out and might be one way reconciling a model of gradual evolution with adaptation to a rapidly changing environment. Gene and genome duplication has been suggested before as a way to explain ‘saltational’ jumps in evolution (53–55). Indeed, the duplication of genes and particularly the duplication of entire genomes immediately creates redundant informational ‘entities’ or ‘modules’ in the genome, offering possibilities for a more drastic change and a wider exploration of genotypic and phenotypic space. While in a constant environment, in which organisms are already fairly well-adapted, we only could observe the further slow optimization of biological systems, a WGD created more drastic changes in certain individual species. These individuals often remained hidden in the population – and usually had a hard time competing with their non-polyploid progenitors, for reasons explained higher and elsewhere (6, 7, 27), but could ‘grab their chance’ upon different conditions or under different contexts. Increased gene pleiotropy and modularity evoked by WGD thus form a fertile substrate allowing evolution to explore more diverse options possibly explaining those bigger jumps in evolution observed. Again, such evolvability might only be successful when the existing environmental conditions have been (seriously) disrupted. When a new environmental ‘equilibrium’ has been reached and the new environmental conditions have stabilized again, the increased evolvability through WGD will become less important in further optimizing the system and on the contrary selection on a more complex system (due to WGD, see higher) might be constrained again (Fig. 4). Considering polyploidy in the evolutionary history of organisms might thus be one additional way of possibly explaining bigger jumps or major transitions in evolution, but certainly needs further investigation.

## Conclusions

Our observations seem to support the two-bladed sword role claimed for WGD in evolution, constituting on the one hand an evolutionary dead end in stable environments, but on the other hand offering opportunities to avoid or reduce the risk of extinction in environmentally unstable and/or drastically changing environments (1, 6). Furthermore, we hope that this work shows the usefulness of our DO-based evolution simulation computational framework in studying the role of WGD in evolution and adaptation, by tracking the evolutionary trajectories of individual artificial genomes in terms of sequence and expression diversification, helping to overcome the traditional limitations of evolution experiments with model organisms.

## Methods

### Description of the Simulation Framework

Below, we provide a summarized description of the different components of our simulation framework, as well as their interactions, focusing on the ones relevant for the current study. A more detailed description of the framework can be found in Yao et al. (25).

### Digital Organism (DO)

Every DO is defined by a genome and a set of i) virtual sensors, which allow DOs to measure different ecological and physiological parameters and ii) actuators, which allow them to interact with each other and with the environment and define the overall behavior of the DO (25) (Fig. 1). For the purposes of this work, we will focus on the actuators replication and movement. However, DOs also have the ability to attack, compete or cooperate with each other and the detail of these functions has been elaborated on previously (24, 25).

### Genes and Genomes

In our framework, the genome of a DO has been inspired by the model previously proposed by Reil (56), and consists of a randomly created string of four digits representing each of the four nucleotides with a total initial size of 100 kb. Genes in the genome are not pre-specified, but identified in the randomly built genome, and typically a genome contains about 150-200 genes (Fig. 2). Up to 1000 different gene products can be encoded by a single genome, each with a unique pre-specified cis-sequence of four digits, identity number and decay rate. Compared to the initial model of Reil, our model makes an explicit distinction among three kinds of genes; i) structural genes, which control every particular behavior of the DO or actuator (e.g., replication or movement), either repressing or promoting it (Fig. *2A*); ii) regulatory genes, encoding transcription factors (TFs), which can specifically activate or repress (50% of each class) the expression of other genes by binding to specific cis-elements in their promoters (Fig. *2B*); and iii) a gene encoding ‘RNA polymerase’, which specifically binds to a ‘TATA’ box (Fig. *2C*).

### Gene Expression

The expression level of a particular gene is calculated based on the TF binding cis-elements found in the promoter of that gene (Fig. 2). When the gene expression level reaches a minimum threshold, the gene is translated into its corresponding gene product. The total amount of gene product (i.e. dosage) corresponds to the expression level of that gene. At every time step during the simulation, the total amount of gene product decays gradually, mimicking protein degradation, with a specific rate that is randomly initialized at the beginning of the simulation. In turn, the decay of gene product is counteracted by the activity of activating TFs on that particular gene. The concentration level (i.e. dosage) of all gene products defines the global GRN of the DO, which is translated into a specific set of actuators (57), which in turn determines the overall behavior of the DO (Fig. 1) (25).

### Signaling Pathways

Signaling pathways are defined as linear equations of sensor inputs. At each time step during the simulation, the value of each kind of sensor input was estimated and used as a variable in the equation. When the result of the equation corresponds to the identity number of a particular gene, its expression is activated (Fig. 1). Linear equations are encoded in the genetic sequence of the genome of a DO, and can change in number and sequence through WGDs and substitutions, respectively. This way, upon initialization of a new DO, a set of 12 TFs is activated (24, 25). Subsequently, at every time step during the simulation, the linear equations compute a new set of identity numbers, increasing the concentration of the corresponding gene product by one unit.

### Food

Represents a source of energy. Each food source represents from 10,000 up to 20,000 energy units. The energy of a food source increased 300 units at every time step until it reached 20,000. At that point, the food source was divided into two food sources, with the newly created food source being randomly placed around the old one. When a DO finds a food source, it assimilates its energy level, and the food source is removed. In our framework, environmental challenges were introduced through the removal of food sources at specific rates and time points.

### Running Simulations

In our evolutionary framework, populations of clonal DOs evolve under fixed WGD and substitution mutation rates, resulting in the evolutionary diversification of the population (Fig. 3). Rather than introducing all polyploids at once in the simulation, we decided to gradually introduce them by setting a fixed WGD rate, which is a situation closer to the real evolutionary process (6, 58). At the beginning of the simulation, the WGD rate was arbitrarily fixed at 40%. When the population size of polyploids reached an equilibrium with that of the non-polyploids (usually in 100 to 200 time steps), the WGD operator was permanently removed. This high rate of WGD as compared with what would be expected in nature was determined by lower WGD rates leading to extremely long computation times until polyploids and non-polyploids reached similarly sized populations. Because our simulations focused less on the initial establishment of polyploids, but more on the potential longer-term advantage of doubling genome content and structure, we decided to start with populations of similar size as the initial condition of the competition. Similarly, at every replication step, a fixed substitution rate (10^−4^ per base) was applied to the offspring genome.

DOs move in a matrix composed of 1000 x 1000 cells looking for food. Every time a DO moves into a neighboring cell in the matrix, its energy level drops by 70 units. DOs were removed from the population when their energy levels dropped below 0. When the energy level of a particular DO reached a threshold of 20,000, the DO was considered to be adapted, and was therefore allowed to replicate (a sign of fitness). However, the final decision of a DO to replicate depends on the relative concentration of the structural gene products involved in promoting and repressing replication. When both concentration levels are equal, the replication has a 50% possibility to happen at that time step.

In order to mimic drastic environmental challenges (for instance as caused by an extinction event), we introduced drastic and/or repeated drops of food during our simulations. In order to simulate different evolutionary scenarios, we applied different food removal patterns. To investigate the interaction between adaptive potential and WGD, we examined the mutational landscape of the genomes across these various evolutionary and environmental contexts. For every DO and at every time step in the simulation, we computed its ploidy and energy level, as well as i) the short-term genetic distance (STGD), defined as the average number of substitutions (per 10 kb) between the individual DO and its direct parent; ii) the long-term genetic distance (LTGD), defined as the average number of substitutions per 10 kb between the individual DO and the initial ancestor at time 0 in the simulation; and iii) the expression distance (ED), defined as the average of the concentrations of all gene products in a given DO compared to the average of the concentrations of all gene products in the previous time step. When the ED differs by more than 30% (arbitrarily defined) between two time steps, we considered the corresponding DO as having an unstable expression pattern. In our simulation, a stable expression pattern (considered to reflect homeostasis) is necessary to maintain a certain ‘phenotype’ and is considered as a sign that the DO has adapted to its environment. Therefore, a sequence of low ED values helped us to identify those DOs that have evolved effective strategies for adaptation.

## ACKNOWLEDGMENTS

Y.V.d.P. acknowledges funding from the European Union Seventh Framework Programme (FP7/2007-2013) under European Research Council Advanced Grant Agreement 322739 – DOUBLEUP. The authors would also like to thank Arthur Zwaenepoel and Eshchar Mizrachi for helpful discussions.

